# Disruption of integrin alpha-5/beta-1-dependent transforming growth factor beta-1 signaling pathway attenuates vessel co-option in colorectal cancer liver metastases

**DOI:** 10.1101/2022.05.24.493291

**Authors:** Miran Rada, Audrey Kapelanski-Lamoureux, Oran Zlotnik, Stephanie Petrillo, Anthoula Lazaris, Peter Metrakos

## Abstract

Colorectal cancer liver metastases (CRCLM) have two major histopathological growth patterns (HGPs) including angiogenic desmoplastic HGP (DHGP) and non-angiogenic replacement HGP (RHGP). The RHGP lesions obtain their blood supply through vessel co-option, where the cancer cells hijack the pre-existing blood vessels of the surrounding liver tissue. Consequently, anti-angiogenic therapies are less efficacious in CRCLM patients with RHGP lesions. Recently, we identified a positive correlation between the expression of Angiopoietin1 (Ang1) and the development of vessel co-opted CRCLM lesions in vivo. However, the mechanisms underlying Ang1 upregulation in vessel co-opting CRCLM lesions are unclear. Herein, we demonstrated that transforming growth factor β1 (TGFβ1) modulates the expression of Ang1 in hepatocytes in vitro. Significantly, pharmaceutical inhibition of integrin alpha-5/beta-1 (ITGα5β1) through ATN-161 impaired TGFβ1-dependent Ang1 expression in vitro and in vivo. Moreover, blocking ITGα5β1 attenuated the formation of vessel co-opting lesions. Furthermore, treatment with ATN-161 significantly improved survival in tumour-bearing mice. Taken together, our results suggest the molecular mechanism of Ang1 upregulation in vessel co-opting CRCLM and targeting this pathway may serve as promising therapeutic strategy to overcome the development of vessel co-option in CRCLM.

## Introduction

Colorectal cancer (CRC) is a leading cause of tumour-related morbidity and mortality worldwide^1^. Approximately, 50% of CRC patients will develop liver metastases (LM), which is the most common cause of mortality^2^. Surgical resection is considered the only curative option resulting in 5 year survival rates of up to 50%^3,4^. However, 80% of colorectal cancer liver metastasis (CRCLM) patients are not resectable^5^. Downsizing with the chemotherapy and targeted therapies such as anti-angiogenic agents (e.g. Bevacizumab) can convert an additional 10–15% of these patients to a resectable state^5–7^.

CRCLMs display two major histopathological growth patterns (HGPs): desmoplastic HGP (DHGP) and replacement HGP (RHGP)^8,9^. The desmoplastic lesions have a desmoplastic layer that separates the cancer cells from the rest of the liver tissue, the cancer cells and hepatocytes do not make contact and they obtain their blood supply via sprouting angiogenesis^9^. In contrast, the cancer cells in replacement lesions make contact with the hepatocytes at the tumour-liver interface and invade the liver parenchyma replacing the hepatocytes which allows them to obtain their blood supply by co-opting pre-existing liver sinusoidal vessels instead of inducing angiogenesis^10–12^. Since anti-angiogenic agents were designed to target only new blood vessel growth, vessel co-option has been implicated as a mechanism of intrinsic and/or acquired resistance to anti-angiogenic therapy in various types of cancer including CRCLM^9,13,14^. In this context, limited responses to anti-angiogenic therapy (bevacizumab) combined with chemotherapy have been observed in CRCLM patients with RHGP lesions compared to patients with DHGP lesions who achieve a better response to the same therapy^9–11,15^. Therefore, deciphering the molecular mechanism(s) vessel co-option and developing therapeutics that would target these mechanism(s) has become of utmost importance.

We previously reported a positive correlation between angiopoietin-1 (Ang1) upregulation in liver parenchyma of CRCLM patients and vessel co-option^15^. However, the mechanisms underlying Ang1 upregulation are unknown. Using an Ang1 knock-out mouse model, we also found that Ang1 deficiency in the liver parenchyma abrogated the development of vessel co-opting lesions through unknown mechanisms^15^. In the current study, we provide mechanistic pathways by which Ang1 upregulated in vessel co-opting CRCLM.

## Results

### Cancer cells promote Ang1 expression in hepatocytes

Recently, we reported the association between Ang1 upregulation in the liver and vessel co-opting CRCLM lesions^15^. It has been reported that TGFβ1 positively correlates with the expression of Ang1 in lung epithelial cells^16^. Importantly, we previously demonstrated upregulation of TGFβ1 in the liver parenchyma of vessel co-opting CRCLM lesions compared to their angiogenic counterparts^17^. Therefore, we hypothesized that TGFβ1 may contribute to Ang1 overexpression in the hepatocytes. To address this question, we exposed IHH hepatocytes to different concentrations (0, 25, 50 and 100 picomolar) of recombinant TGFβ1 for 24 hours (Figure 1a). Indeed, exposing hepatocytes to recombinant TGFβ1 promoted the expression of Ang1. IHH cells are immortalized human hepatocytes that retained several differentiated features of normal hepatocytes^18–20^.

**Figure 1.**
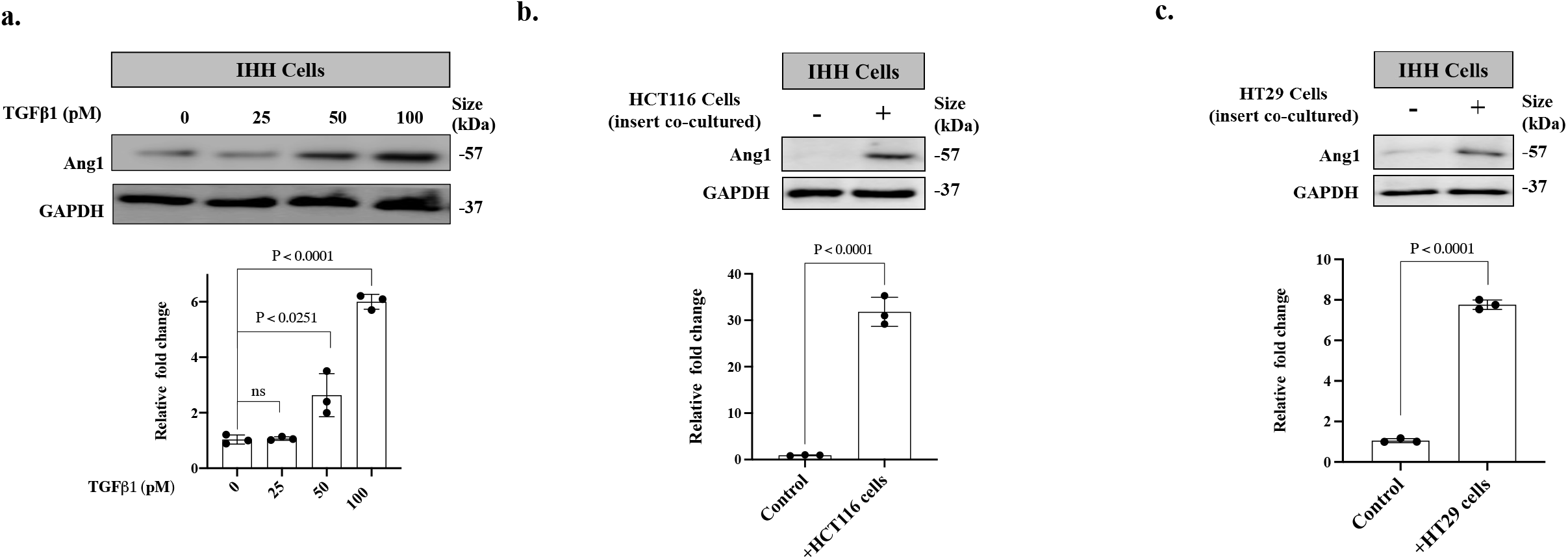
Cancer cells induce Ang1 expression in hepatocytes through TGFβ1. **a**. Western blot of Ang1 expression in IHH hepatocytes in the presence of various concentration (0, 25, 50, 100 picomolar) of recombinant TGFβ1 (Top panel). Bottom panel shows the intensity of Ang1 bands that quantified and normalized against GAPDH using ImageJ and represented as a fold change (n=3). **b**. Upper panels represent immunoblotting of Ang1 expression in human hepatocytes (IHH cells) co-cultured with HCT116, HT29 colorectal cancer cells respectively. Bottom panel shows the intensity of Ang1 bands that quantified and normalized against GAPDH using ImageJ and represented as a fold change (n=3). Data are presented as the mean ± SD.

Our previous publications suggested higher levels of TGFβ1 in the conditioned media of co-cultured IHH hepatocytes with colorectal cancer cells in comparison to the conditioned media of the same cells individually, which is incited by higher expression levels of TGFβ1 in the co-cultured hepatocytes and cancer cells^17,21^. Therefore, we questioned whether co-culturing IHH hepatocytes with cancer cells induce the expression of Ang1 through TGFβ1. Our immunoblotting data suggested overexpression of Ang1 in the co-cultured hepatocytes compared to their control (Figure 1b and 1c).

To confirm the co-localization of Ang1 and TGFβ1 in CRCLM specimens, we performed co-immunofluorescence staining on chemonaïve human sections using anti-TGFβ1 and anti-Ang1 antibodies. Our results demonstrated overexpression and co-localization of both proteins in vessel co-option lesions (Figure 2a). Notably, both TGFβ1and Ang1 staining were more pronounced at the tumour-liver interface where the hepatocytes of the normal adjacent liver and cancer cells are in very close proximity. Significantly, some cancer cells in the peripheral tumour were positively stained for TGFβ1 and Ang1. To identify unequivocally in which tissue compartment the expression of TGFβ1 and Ang1 were upregulated, we performed fluorescence in situ hybridization (FISH) assay for TGFβ1 and Ang1 combined with cancer cell specific (anti Cytokeratin 20 antibody) staining. The RNA levels of both TGFβ1 and Ang1 were selectively upregulated in the adjacent normal liver parenchyma (most likely hepatocytes) in CRCLM lesions and not in the cancer cells (Figure 2b). Consequently, any positive staining for TGFβ1 and Ang1 in adjacent cancer cells is most likely due to the uptake of the secreted TGFβ1 and Ang1 from the liver parenchyma (hepatocytes) rather than their expression. Collectively, our data proposed the interaction between cancer cells and hepatocytes as a key factor for Ang1 overexpression, which is mediated by TGFβ1.

**Figure 2.**
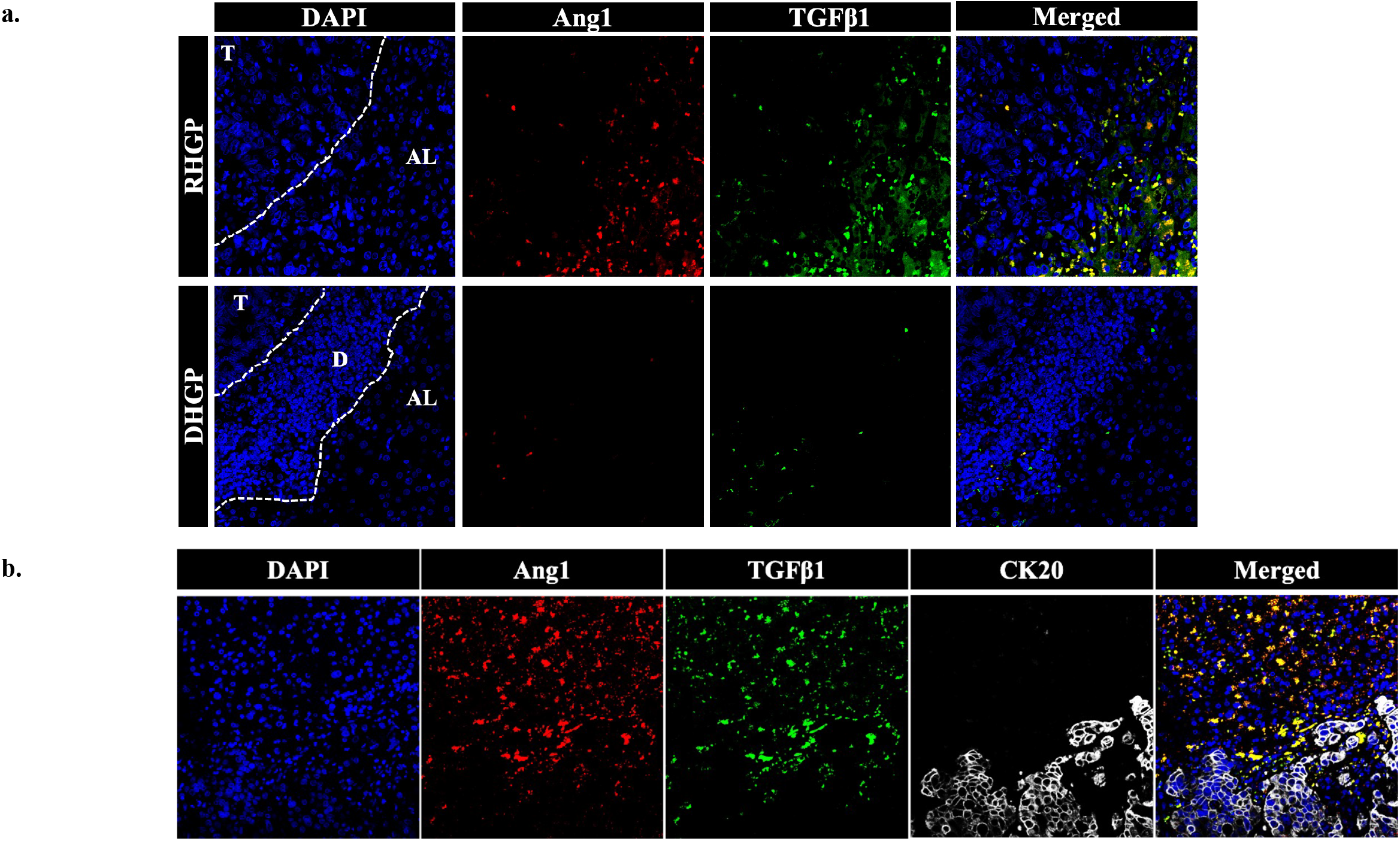
Hepatocytes are the source of TGFβ1 and Ang1 expression. **a**. Immunofluorescence staining of chemonaïve CRCLM lesions showing Ang1 (red) and TGFβ1 (green). AL: Adjacent liver, D: Desmoplastic ring, T: Tumour **b**. Fluorescence in situ hybridization (FISH) for Ang1 (red) and TGFβ1 (green) mRNA expression in chemonaïve CRCLM lesions overlapped with CK20 (cytokeratin 20, white) antibody.

### TGFβ1-dependent Ang1 expression in hepatocytes is mediated by ITGα5β1

Integrins comprise a family of heterodimeric extracellular matrix receptors that mediate the interaction between cells and various microenvironments^22-24^. Importantly, integrin subunits have been reported as a mediator of TGFβ1 signalling cascade in various tumours^25,26^. Hence, we decided to explore the expression levels of various integrin subunits in chemonaïve CRCLM lesions including integrin alpha-5 (ITGα5) and integrin beta-1 (ITGβ1). Our data suggested dramatic upregulation of both ITGα5 (Figure 3a) and ITGβ1 (Figure 3b) in vessel co-opting lesions compared to angiogenic lesions. Subsequently, we sought to assess the role of ITGα5β1 in TGFβ1-dependent Ang1 overexpression in hepatocytes. We treated IHH hepatocytes with a pharmacological inhibitor against ITGα5β1 (ATN-161)^27,28^ upon exposure to recombinant TGFβ1. As shown in Figure 3c, the effect of TGFβ1 was dramatically weakened in the presence of 8.0μM ATN-161.

**Figure 3.**
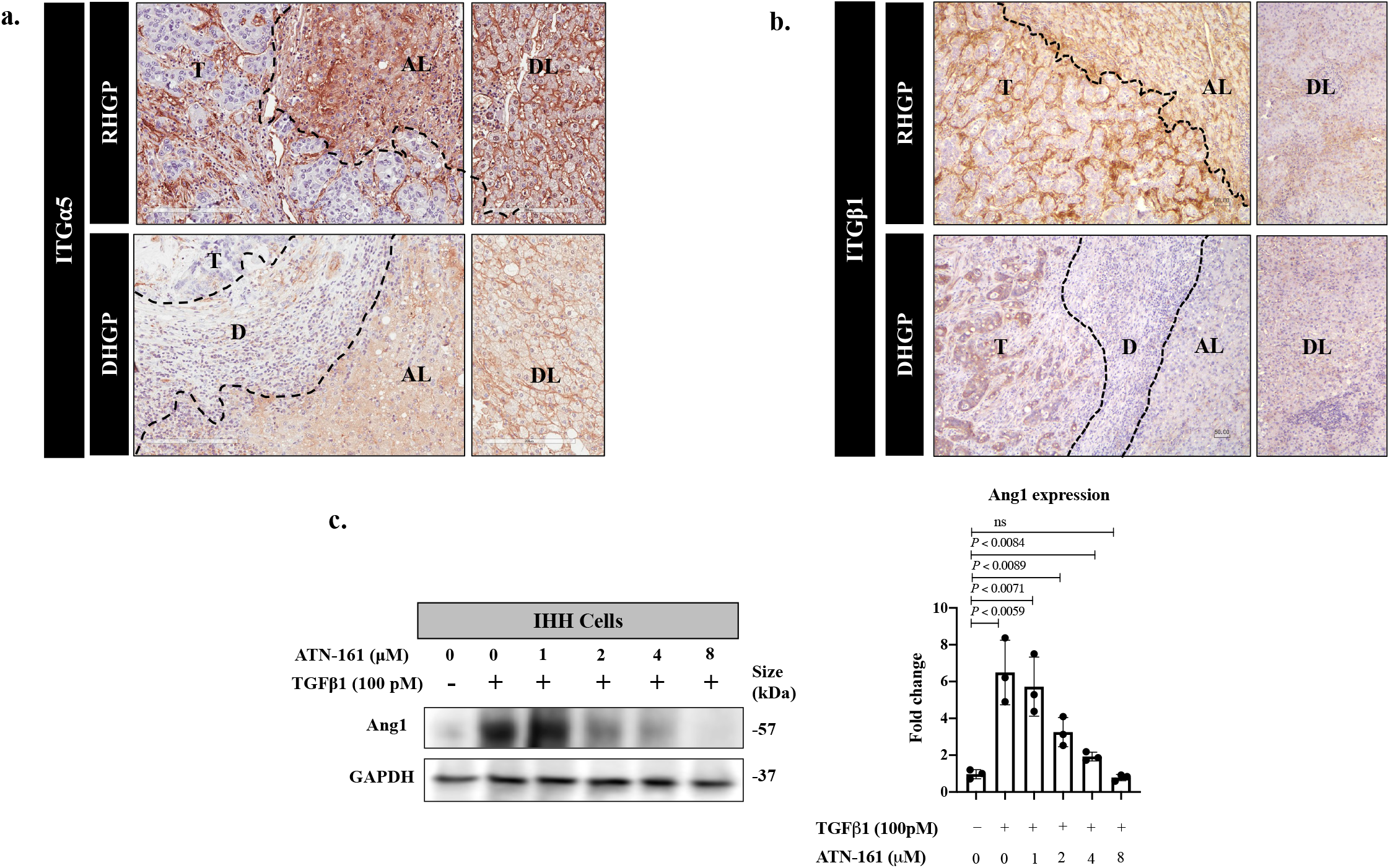
ITGα5β1 mediates TGFβ1-dependent Ang1 expression in hepatocytes. **a** and **b**. Representative immunohistochemistry images of ITGα5 and ITGβ1 in chemonaïve CRCLM lesions respectively (n= 5 RHGP, n=5 DHGP). AL: Adjacent liver, D: Desmoplastic ring, DL: Distal liver, T: Tumour. **c**. Left panel represents immunoblotting of Ang1 expression in human hepatocytes (IHH cells) upon treatment with recombinant TGFβ1 in the presence or absence of ITGα5β1 inhibitor (ATN-161). The right panel shows the intensity of Ang1 bands that were quantified and normalized against GAPDH using ImageJ and represented as a fold change (n=3). Ns= no significant. Data are presented as the mean ± SD.

To further confirm the role of ITGα5β1 in TGFβ1 signaling, we used ATN-161 in vivo. As shown in 4a, we injected MC38 cancer cells intrasplenically into wild-type C57B/6 mice (n=6) followed by treatment with ATN-161 (n=3) or placebo (n=3). The treatment was performed in 5 doses (200μM every 2 days). Intriguingly, blocking ITGα5β1 significantly extended the survival of the mice (Figure 4b). Moreover, it also reduced the size (Figure 4c) and the number of the generated liver metastatic lesions (Figure 4d). More importantly, treating mice with ATN-161 induced the formation of angiogenic DHGP lesions (Figure 4e). Indeed, the expression levels of Ang1 in the adjacent liver were also decreased upon inhibition of ITGα5β1 (Figure 4f). Together, our results propose that ITGα5β1 is an essential molecule for the formation of vessel co-opting CRCLM lesions.

**Figure 4.**
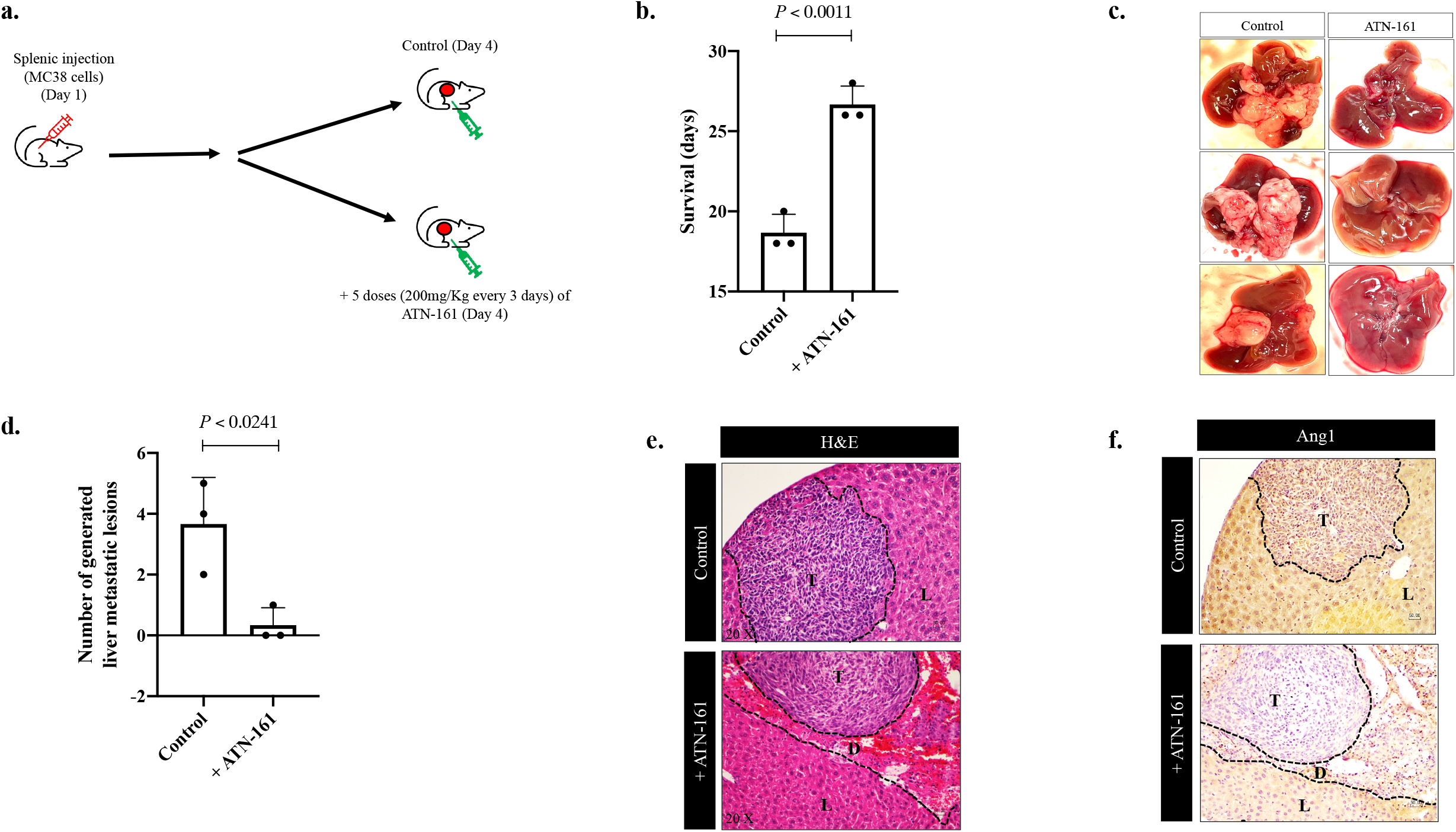
Blocking ITGα5β1 attenuates the formation of vessel co-opting lesions. **a**. Schematic representation of the experimental design. **b**. Represents the survival (days) of the tumour-bearing mice. **c**. Shows the liver of the tumour-bearing mice in the presence or absence of ITGα5β1 inhibitor (ATN-161). **d**. Shows the number of the generated liver metastatic tumours. **e**. Represents hematoxylin-eosin (H&E) staining of the liver metastatic sections. **f**. Represents immunohistochemical staining of the generated tumours using Ang1 antibody. D: Desmoplastic ring, L: Liver, T: Tumour. Data are presented as the mean ± SD.

## Discussion

During the last decades, angiogenesis has been regarded as the most important process by which cancer cells vascularize themselves, while alternative vascularization mechanisms were ignorored^29^. Alternative vascularization mechanisms include vessel co-option^14^, increased pericyte coverage^30^, vasculogenic mimicry^31^, lysosomal sequestration^32^ and Glycosylation-dependent angiogenesis^33^. However, vessel co-option has emerged as the main alternative vascularization pathway that mediates the failure of anti-angiogenic treatment in CRCLMs^9^.

In vessel co-opted CRCLM tumours, the cancer cells meet their metabolic demands without the generation of new vessels^9,10^. The cancer cells migrate and infiltrate the surrounding liver tissue space between pre-existing vessels, ultimately leading to the incorporation of pre-existing vessels into the tumour^8,12,15^.

Ang1 has been recognised as bona fide ligand of the Tie2 receptor, which serves an essential role in vascular development and angiogenesis^34^. Recently, we found Ang1 as a mediator of vessel co-option formation in CRCLM^15^. Importantly, the function of Ang1 in vessel co-opting lesions depends on cancer cell motility^35^. Accordingly, Ang1 induces the expression of ARP2/3 via Tie2-PI3K/AKT pathway^35^. However, the upstream signaling of Ang1 expression in CRCLM is poorly understood. Herein, we suggested a mechanistic pathway that involved in Ang1 overexpression in vessel co-opting tumours, which is regulated by TGFβ1.

TGFβ1 is known to play dichotomous roles in tumour angiogenesis. In this context, TGFβ1 has been reported as an inducer of angiogenesis, which promotes the expression of vascular endothelial growth factor A (VEGFA) and/or other angiogenic factors in cancer cells^36,37^. In contrast, Geng et al.^38^ have shown that TGFβ1 signalling pathway blocks angiogenesis in metastatic colon cancer by increasing VEGFA ubiquitination and degradation. Our results corroborated that expression of TGFβ1 is associated with non-angiogenic (vessel co-opting) tumours. Various investigations have proposed cancer cells as the main source of TGFβ1 expression and secretion in tumour^39-41^, while our current study identified hepatocytes as the primary source of TGFβ1 expression in CRCLM. Different TGFβ1 receptors have been found to facilitate TGFβ1 interaction with tumour cells including ITGα5β1^42,43^. In agreement with these studies, we observed attenuation of TGFβ1 function in hepatocytes by blocking ITGα5β1 through (ATN-161) in vitro and in vivo. It is worth mentioning that ATN-161 has been shown to have antitumor activity in a number of different preclinical models^44^. Various studies suggested that ATN-161 has a potential therapeutic index. Therefore, the effect of ATN-161 has been evaluated in different clinical trials. Testing of ATN-161 in Phase I clinical trial has revealed that this compound has anti-tumour activity and is well tolerated in patients with solid tumours^45^. Therefore, the current study presents new insights into the targeting TGFβ1 receptors in tumour treatment.

In conclusion, the present study proposed the mechanistic pathways for Ang1 upregulation in vessel co-opting CRCLM tumours (Figure 5). In addition, this study provides evidence that ITGα5β1 has a notable role in the development of vessel co-opting lesions, which could be used as potential novel therapeutic targets in future to treat vessel co-opted CRCLM lesions.

**Figure 5.**
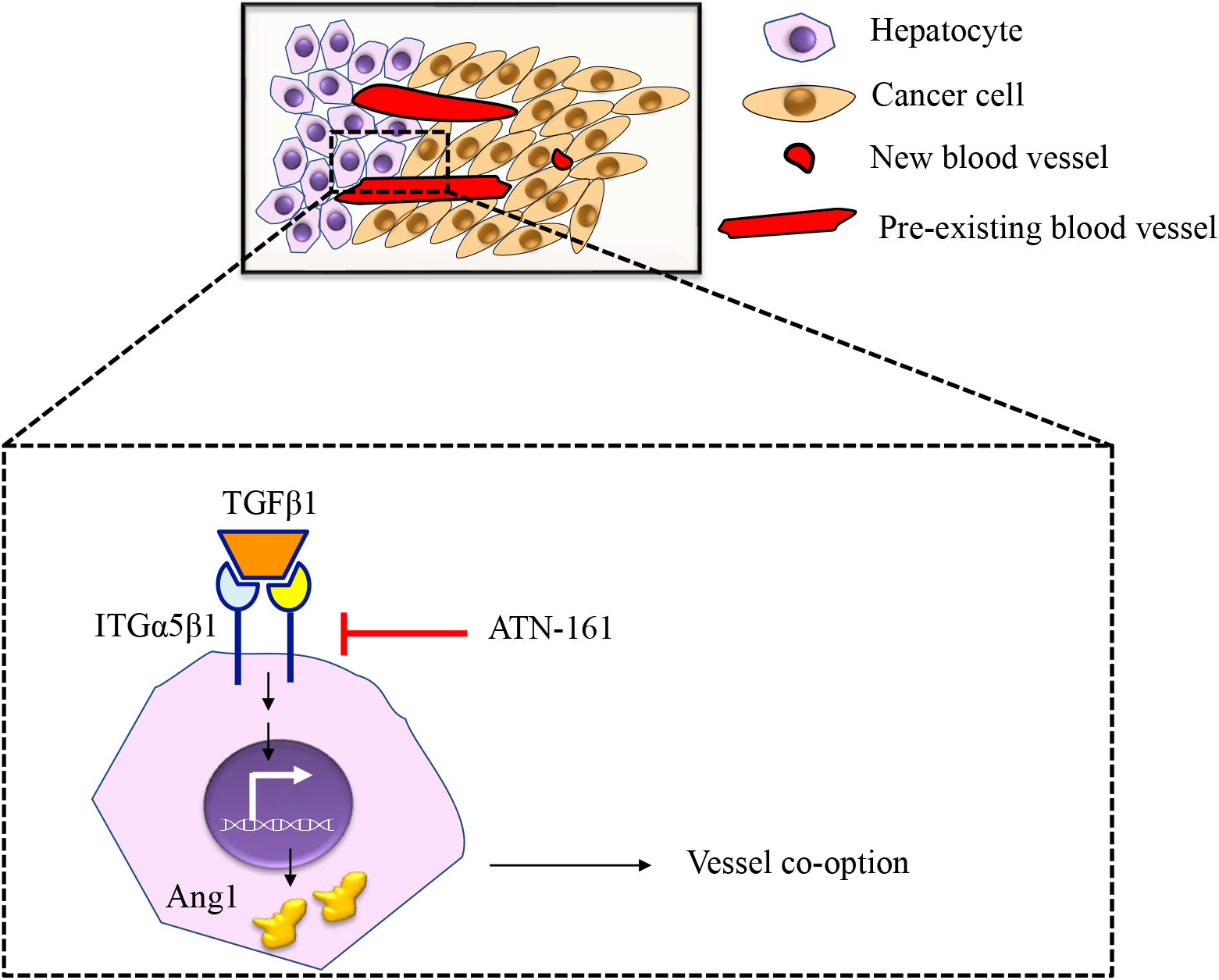
The mechanism of Ang1 upregulation in vessel co-opting CRCLM. Schematic representation of key findings in the current study. The secreted TGFβ1 by adjacent hepatocytes contributes to Ang1 expression in the hepatocytes through ITGα5β1. The secreted Ang1 induces vessel co-option.

## Materials and methods

### Patient samples

The study was conducted in accordance with the guidelines approved by McGill University Health Centre (MUHC) Institutional Review Board (IRB). Informed consent was obtained from all patients through the McGill University Health Centre Liver Disease Biobank (MUHC-LDB). Surgical specimens were procured and released to the Biobank immediately after the pathologist’s confirmation of carcinoma and surgical margins.

### Cell cultures

HT29 colorectal cancer cell line was a gift kindly supplied by Dr Alex Gregorieff (Cancer Research Program, McGill University). HCT116 was kindly provided by Dr Daniela Quail (Rosalind and Morris Goodman Cancer Research Centre, McGill University). IHH cells were a generous gift from Dr Nabil G. Seidah at Montreal Clinical Research Institute (IRCM). The cells were cultured in DMEM (Wisent Inc., #319-005-CL) supplemented with 10% FBS (Wisent Inc., #085-150) and 1× penicillin/streptomycin (Wisent Inc., 450-201-EL). All cells were cultured at 37 °C with 5% CO2. To block the ITGα5β1 pathway, we treated cells ATN-161 (Tocris, #6058) for 24 hours using concentrations indicated in the figures.

### Co-culture, treating cells with recombinant TGFβ1

Before the experiments with recombinant TGFβ1 (Peprotech, #100-21), the cells were seeded in DMEM (Wisent Inc., #319-005-CL) supplemented with 10% FBS (Wisent Inc., #085-150) and 1× penicillin/streptomycin (Wisent Inc., 450-201-EL) for 24 hours. The next day, the conditioned media was aspirated, and the cells were washed with PBS (Wisent Inc., # 311-010-CL) twice. Then, serum-free DMEM supplemented with either TGFβ1 or Ang1 was added, and the cells were incubated for 24 h at 37 °C.

Co-culturing was conducted using 6-well inserts (Falcon, #353090) and companion plates (Falcon #353502). The cells were cultured with DMEM (Wisent Inc., #319-005-CL) supplemented with 10% FBS (Wisent Inc., #085-150) and 1× penicillin/streptomycin (Wisent Inc., 450-201-EL) overnight. The next day, the media was removed, and the cells were washed twice with PBS (Wisent Inc., # 311-010-CL). New serum-free DMEM (Wisent Inc., #319-005-CL) was added and incubated at 37 °C for 24 hours.

### Immunoblotting

To prepare lysate from the cultured cells, the medium was removed, and cells were washed once with 1x PBS, trypsinized, collected and kept on ice. Cells were ruptured by passing through a syringe 10 times and centrifuged for 10 minutes at 5000 rpm. The supernatant was transferred into 1.5mL microcentrifuge tubes, and protein concentrations were determined using BCA Protein Assay Kit (Thermo Scientific, #23225). 5–10 μg of total protein per sample were subjected to 10-12% SDS-PAGE and transferred to Immobilon-E membranes (Millipore, #IEVH85R). The blots were developed using Pierce ECL Western Blotting Substrate (Thermo Scientific, #32106) and imaged with ImageQuant LAS4000 (GE Healthcare BioScience). The intensity of the bands was assessed using ImageJ (NIH, Bethesda, MD) software^46-50^.

The following primary antibodies were used: GAPDH 1:2000 (Abcam, # ab9485) and Ang1 1:1000 (Abcam, #ab102015).

### Immunohistochemical staining

Formalin-fixed paraffin-embedded (FFPE) CRCLM resected blocks, provided through the MUHC-LDB, were used for this study. Serial sections 4 mm thick were cut from each FFPE block, mounted on charged glass slides (Fisher Scientific, #12-550-15) and baked at 37°C overnight. Prior to staining, the slides were baked at 60°C for 1 hr as well. Hematoxylin and eosin (H&E)-stained sections were prepared from all cases for an initial histopathological assessment. The sections were deparaffinized with xylene (Leica, #3803665) followed by hydration with graded concentrations of ethanol (Comalc, #P016EAAN) and then with distilled water. Samples were subjected to antigen retrieval (Sodium Citrate 10mM, pH=6.0) followed by washing with PBS and incubation in hydrogen peroxide (Dako, #S2003) to inhibit endogenous peroxidase. The tissue sections were blocked with 1 % goat serum and incubated with the indicated primary antibody in 1 % goat serum overnight at 4°C. After washing, the sections were incubated with secondary antibody (Dako, Anti-Mouse: #K4001; Anti-Rabbit: #K4003) for 1 h at room temperature and positive signals were visualized with the diaminobenzidine (DAB) substrate (Dako, #K3468). The following primary antibodies were used: ITGα5 1:400 (Cell Signaling, #4705) and ITGβ1 1:200 (Cell Signaling, #9699).

All slides were scanned at 20× magnification using the Aperio AT Turbo system. Images were viewed using the Aperio ImageScope ver.11.2.0.780 software program for scoring analysis and assessment of signals.

### Immunofluorescence staining

Formalin-fixed paraffin-embedded (FFPE) human CRCLM resected blocks were deparaffinized with xylene followed by hydration with graded concentrations of ethanol and then with distilled water. Samples were subjected to antigen retrieval (Sodium Citrate 10mM, pH=6.0) followed by washing with PBS and incubation in hydrogen peroxide (Dako, #S2003) to inhibit endogenous peroxidase. The tissue sections were blocked with 1 % goat serum and incubated with the indicated primary antibody in 1 % goat serum overnight at 4°C. After washing, the sections were incubated with secondary antibody 1:1000 (Alexa Flour 594 gaot anti-rabbit IgG and Alexa Flour 488 goat anti-mouse IgG (Invitrogen #A11037 and #A10680 respectively)) for 1 h at room temperature followed by washing thrice. The sections were incubated with 4′,6-Diamidino-2-Phenylindole, Dihydrochloride DAPI 1:1000 (Thermo Fisher Scientific, D1306) in PBS for 10 minutes at room temperature. Prior to mounting under cover glasses, 1-2 drops of ProLong Gold Antifade Mountant (Thermo Fisher Scientific, P36934) were added to each section.

The following primary antibodies were used: TGFβ1 1:1500 (Abcam, # ab27969), Ang1 1:1500 (Abcam, # ab102015) and Cytokeratin 20 1:100 (abcam, #ab76126).

### Fluorescence in situ hybridization (FISH)

To identify TGFβ1 and Ang1 co-expression in CRCLM lesions fluorescence in situ hybridization was performed according to the manufacturer’s recommendations^15^ using RNAscope Probe-Hs-TGFβ1, labelled with Alexa 488 nm fluorescent dye. Briefly, formalin-fixed paraffin-embedded (FFPE) human CRCLM sections (4 μm) were baked for 1h at 60°C. The sections were deparaffinized through successive baths of xylene (100%), ethanol (95%) and then distilled water. After drying, the slides were incubated for 10 min with RNAscope Hydrogen Peroxide at room temperature followed by washing with distilled water. Then, target retrieval was conducted by incubating the slides with RNAscope 1X Target Retrieval Reagents in a steamer for 20 min. The sections were incubated with ethanol for 3 min, dried, and incubated with RNAscope Protease Plus and incubate at 40°C for 30 min. The slides were washed with distilled water, dried, and hybridization was carried out with RNAscope Probe-Hs-TGFβ1-C2 (ACDBIO, # 400881-C2) and RNAscope® Probe-Hs-ANGPT1 (ACDBIO, # 482901) diluted in a Blank Probe – C1 (ACDBIO, #300041) and incubated in in HybEZ™ Oven (ACDBIO, #321710) at 40°C for 2 hours. The slides were then incubated with SSC buffer (Sigma-Aldrich/MLS, # S6639-1L) overnight at room temperature. The following day, the slides were washed and incubated at 40°C with RNAscope Multiplex FL V2 AMP-1 (ACDBIO, #323110) for 30 min, RNAscope Multiplex FL V2 AMP-2 for 30 min and RNAscope Multiplex FL V2 AMP-3 for 15 min. After washing in 1X PBST, the sections were incubated with RNAscope Multiplex FL v2 HRP-C1 (ACDBIO, #323110) for 15 min at 40°C. The dye was prepared by diluting Opal 520 Reagent in RNAscope Multiplex TSA Buffer 1:1500 (ACDBIO, #322809) and added to the sections for 30 min at 40°C followed by incubation with RNAscope® Multiplex FL v2 HRP blocker for 15 min at 40°C. Next, we incubated the slides in 1% BSA for 30 min at room temperature and staining was performed with Cytokeratin 20 1:100 (abcam, #ab76126) following the abovementioned Immunofluorescence staining protocol. The sections were mounted under cover slips using ProLong Gold Antifade Mountant (Thermo Fisher Scientific, P36934) and visualized with Zeiss LSM780 confocal microscope and Zen software (Zeiss International, Oberkochen, Germany).

### Xenograft experiments

Mouse experiments were performed as previously described^15,23^. The mice were randomly assigned to each group. Colorectal cancer liver metastases were generated in 4-week to 6-weeks old C57B/6 mice by intrasplenic injection of 50 μL of PBS (Wisent Inc., #311010-CL) containing 1 ×10^6^ MC38 cancer cells followed by splenectomy 1 min after injection. On day 4, the treatment started with ATN-161 (5 doses: 200μM every 2 days). The mice were euthanized once their health deteriorated. The experiment was terminated on day 28. The liver was collected and fixed in 10% buffered neutral formalin and embedded in paraffin. Hematoxylin and eosin (H&E)-stained sections were prepared from all samples for an initial histopathological assessment.

The mice were housed in facilities managed by the McGill University Animal Resources Centre. All animal experiments were conducted under a McGill University approved Animal Use Protocol in accordance with guidelines established by the Canadian Council on Animal Care.

### Statistical analysis

Statistical analysis was performed with a two-tailed Student’s t-test using GraphPad Prism software version 8.0 (GraphPad Software, La Jolla, CA, USA) and Excel software. Unpaired t-test was applied to compare means of two groups. Data presented as mean ± standard deviation. Unpaired Student’s t-test was applied to compare the means of two groups. The association between the two categorical groups in xenograft experiments was assessed with the Chi-square test. P-values of <0.05 were considered to be significant.

## Acknowledgments

We thank Dr Alex Alex Gregorieff (Cancer Research Program, McGill University), Dr Daniela Quail (Rosalind and Morris Goodman Cancer Research Centre, McGill University) and Dr Nabil Seidah (Montreal Clinical Research Institute) for providing cell lines. We acknowledge Shaida Ouladan for her support with FISH experiments. We also thank RI-MUHC Liver Disease Biobank for providing human samples. We especially thank our patients who are an inspiration, they give selflessly so that others, total strangers, will one day benefit. This work was supported by RI-MUHC award, Dana Massaro and Ken Verdoni Liver Metastases Research Fellowship.

## Author contributions

M.R., A.L. and P.M. co-conceived the study. M.R. executed the experiments. M.R. performed immunohistochemistry, immunofluorescence, FISH, cell culture, immunoblotting, and mice treatment. A.K.L. assisted in cell culturing and immunoblotting. A.K.L. and O.Z. conducted the splenic injection. S.P. collected and prepared human samples. M.R wrote the manuscript. All authors provided critical reviews and approved the manuscript.

